# Toward single cell tattoos: Biotransfer printing of lithographic gold nanopatterns on live cells

**DOI:** 10.1101/2023.05.28.542285

**Authors:** Kam Sang Kwok, Yi Zuo, Soo Jin Choi, Gayatri J. Pahapale, Luo Gu, David H. Gracias

## Abstract

Lithographic nanopatterning techniques like photolithography, electron-beam lithography, and nanoimprint lithography (NIL) have revolutionized modern-day electronics and optics. Yet, their application for creating nano-bio interfaces is limited by the cytotoxic and two-dimensional nature of conventional fabrication methods. Here, we present a biocompatible and cost-effective transfer process that leverages (a) NIL to define sub-300 nm gold (Au) nanopattern arrays, (b) amine functionalization of Au to transfer the NIL-arrays from a rigid substrate to a soft transfer layer, (c) alginate hydrogel as a flexible, degradable transfer layer, and (d) gelatin conjugation of the Au NIL-arrays to achieve conformal contact with live cells. We demonstrate biotransfer printing of the Au NIL-arrays on rat brains and live cells with high pattern fidelity and cell viability and observed differences in cell migration on the Au NIL-dot and NIL-wire printed hydrogels. We anticipate that this nanolithography-compatible biotransfer printing method could advance bionics, biosensing, and biohybrid tissue interfaces.

**TOC Figure:** 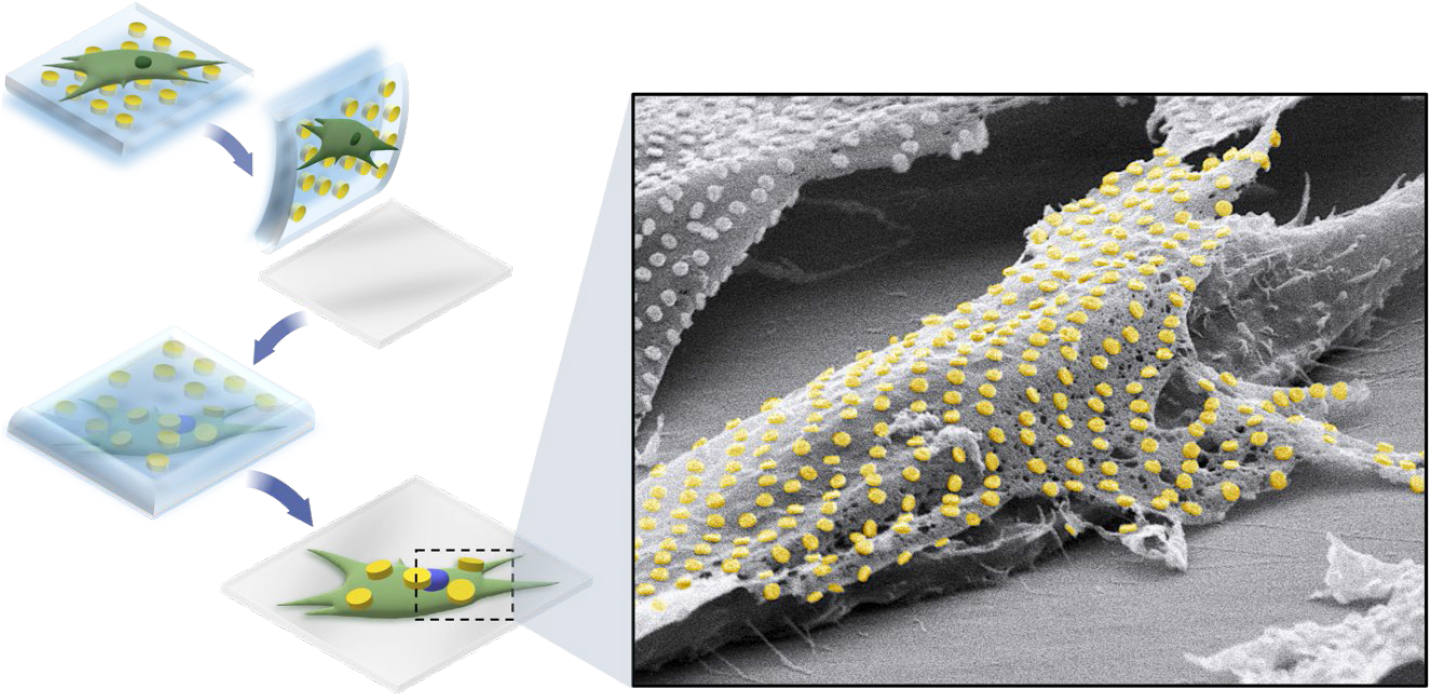

---

Engineers have long sought to merge nanoelectronics, nanophotonics, and stimuli-responsive materials with the human body across length scales of organs to single cells.^1,2^ To create smart devices tailored to the soft, dynamic, and three-dimensional (3D) surfaces of biological systems, it is necessary to establish methods that can reliably integrate well-defined nanopatterns, such as electrode arrays, antennas, and circuits, onto living cells and tissues. In the last few decades, advances in very large-scale integration (VLSI) and micro-electromechanical systems (MEMS) have enabled the fabrication of sophisticated devices like transistors, integrated circuits, and sensors with exquisite nanoscale resolution. More recently, the assembly of materials and devices on flexible substrates that can mold to curvilinear surfaces has been achieved via laser printing,^3–5^ 3D printing,^6^ micro pick-and-place systems,^7^ and self-assembly.^8^ These top-down processes, however, often utilize harsh chemicals, high temperatures, or vacuum techniques that are incompatible with living cells, tissues, and soft, aqueous materials.

To address this challenge, researchers have explored alternative approaches to creating biological interfaces, such as depositing force-mediating nanoparticles on cells or 3D bioprinting composite formulations of nanomaterials and cells.^9–11^ However, these biocompatible techniques often possess limited throughput and resolution, especially at sub-micron length scales. Yet others have shown that living cells can internalize microstructures, such as radio-frequency identification (RFID),^12^ force and pressure sensors,^13,14^ barcodes,^15^ magnetic antennas,^16^ and microrobots.^17^ Indeed, these achievements demonstrate the possibility of interfacing various materials with live cells and tissues. However, a lithography-based technique for systematically integrating nanomaterials onto live cells with high spatial resolution and yield has yet to be realized.

Nanotransfer printing (nTP) offers a high throughput approach to printing large-area arrays of nanopatterns on unconventional 3D substrates,^18^ such as polymers,^19^ elastomers,^20^ and hydrogels.^21^ For instance, Jeong *et al*. used a solvent-assisted nTP technique to print arrays of plasmonic silver nanowires on soft contact lenses for enhanced Raman signals, which enabled glucose detection at low concentrations.^22^ Similarly, Ko *et al*. printed Au nanowires on hyaluronic acid film to develop smart contact lenses capable of treating Irlen syndrome.^21^ While these nTP techniques are capable of printing large-area nanopatterns on flexible substrates in parallel, they require organic solvents (e.g. toluene, acetone), high pressure (e.g. 3 bar), or high temperatures (e.g. 45-100°C)—all of which are highly unfavorable conditions for living systems.

Here, we present a hybrid nTP process that can bond lithographically defined gold (Au) nanopatterns to live cells in physiological conditions. Our approach involves three main steps: 1) conventional thermal nanoimprint lithography (NIL) and subsequent transfer onto glass coverslips to obtain arrays of Au nanodots and nanowires, 2) amine functionalization of the Au NIL-arrays followed by alginate hydrogel casting to delaminate the Au NIL-arrays from the glass coverslip, and 3) chemical conjugation of the Au NIL-arrays with gelatin to assist transfer onto tissue or living cells followed by the dissociation of the alginate hydrogel with ethylenediaminetetraacetic acid (EDTA). In this study, we show that our approach can reliably transfer 8 by 8-mm arrays of Au nanodots (250 nm diameter) and nanowires (300 nm width) created by NIL to soft and flexible alginate hydrogels. We observed pattern-specific cell migration on the Au NIL-array printed hydrogels and optimized alginate hydrogel dissolution with EDTA to maintain high cell viability. After dissociating the alginate hydrogel transfer layer, we observed that the Au NIL-arrays bonded to individual fibroblast cells. Overall, this approach offers a versatile strategy for seamless, tattoo-esque integration of NIL-patterns and arrays with live cells and tissues.

In the first step (Figure S1), we spin-coated a sacrificial layer of polydimethylglutarimide (PMGI, SF6) on a silicon (Si) wafer. We created 8 by 8-mm arrays of Au nanodots and nanowires on the wafer by thermal NIL and thermal evaporation of 5 nm of chromium (Cr) as an adhesion layer and 50 nm of Au. We then spin-coated 200 nm of polymethyl methacrylate (PMMA, A4) as a carrier film on the nanopatterned Si wafer. We released the Au NIL-arrays from the substrate by floating the wafer on top of a positive photoresist developer (MF26A) to dissolve the PMGI sacrificial layer. Afterward, we displaced the photoresist developer with deionized water to rinse the film and subsequently with Cr etchant (Chromium Cermet Etchant TFE) to remove the Cr. After repeating the rinsing step with water, we picked up the film carefully with a glass coverslip. The choice of glass coverslips as the substrate enables efficient transfer of the Au NIL-arrays to the alginate hydrogel in the second step, since Au has relatively poor adhesion to SiO_2_.^23^ Finally, we etched the PMMA film in oxygen plasma to obtain Au NIL-arrays on the glass coverslip. The NIL-arrays can be transferred onto glass coverslips with high fidelity, and it is noteworthy that such patterns can also be transferred onto rigid 3D shapes, such as the microparticle shown in Figure S2.

The transfer of the Au NIL-arrays to live cells and tissues requires additional criteria to be met, including flexibility, physical integrity, compatibility with cell culture media, and appropriately designed relative adhesion. Specially designed hydrogels are an alternative to rigid substrates and can also act as a sacrificial layer by reverse gelation. Alginate is widely used for cell culture and tissue engineering applications due to its biocompatibility and tunable, tissue-mimetic mechanical properties.^24,25^ Therefore, we selected alginate hydrogel as an intermediary substrate to delaminate the Au NIL-arrays from the rigid glass coverslip and affix them to cell sheets and brain tissues. A key requirement for the hydrogel assisted transfer of the Au NIL-arrays in the second step is that the adhesion between the Au NIL-array and the alginate hydrogel is greater than the adhesion between the Au NIL-array and the underlying glass substrate.

We investigated the effect of surface functionalization of Au on the adhesion of the Au NIL-array to the alginate hydrogel using self-assembled monolayers of either 3-mercaptopropionic acid (3-MPA) or cysteamine. Both molecules have a thiol group that can bind to Au, but 3-MPA contains a negatively charged carboxylic acid end group in its deprotonated form,^26^ while cysteamine contains a positively charged amine end group in its protonated form.^27,28^ We immersed the Au NIL-arrays in ethanol solutions containing 0.26 mM of either 3-MPA or cysteamine for an hour. We then intermixed 0.5 ml of 2.5 wt% alginate solution with 125 μl of 25 mM calcium sulfate and cast the resulting alginate hydrogel on the Au NIL-arrays. After allowing the solution to gel for 45 minutes under a glass slide, we gently peeled off the alginate hydrogel containing the Au NIL-array from the glass coverslip and placed it pattern-side up for further characterization (Figure 1). We observed that the transfer yield of the cysteamine-functionalized Au NIL-array was approximately twice that of the 3-MPA-functionalized Au NIL-array (Figure S3). We attribute this difference to favorable electrostatic forces between the positively charged end groups of the cysteamine molecules and the negatively charged carboxyl groups of alginate.^29^ Additionally, the SEM images (Figure 1c-d) show that this transfer process can print both NIL-dots and NIL-wires with high fidelity.

**Figure 1.**
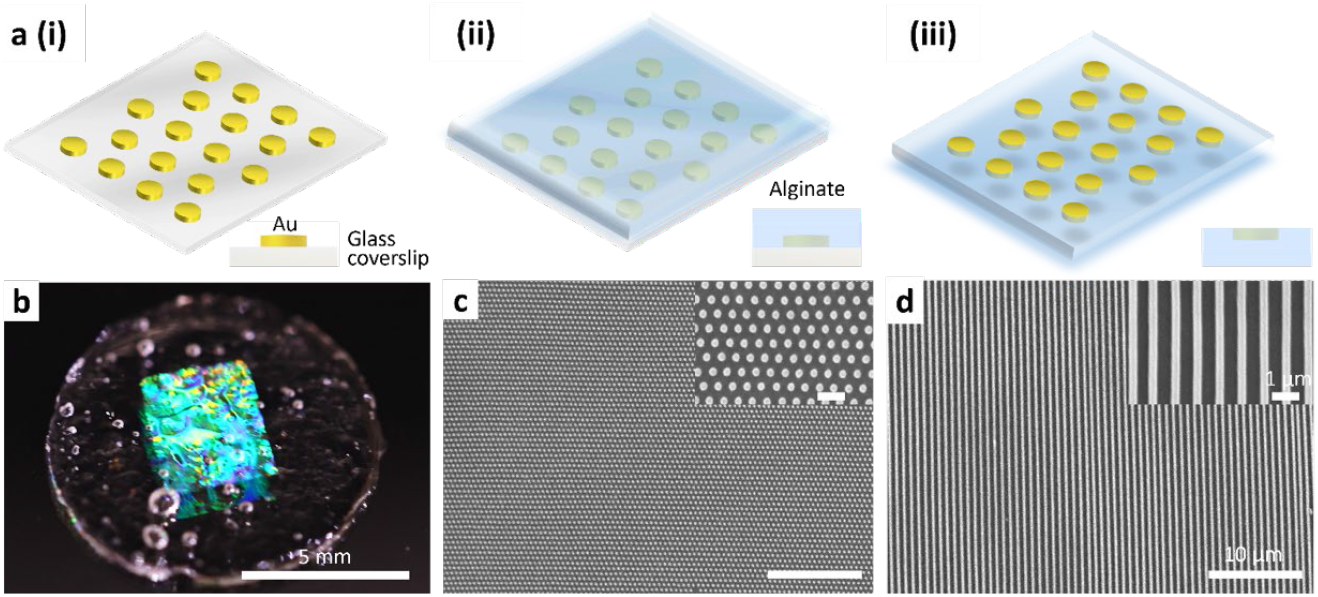
Schematic illustration and microscopy images showing the transfer process of the Au NIL-array to the alginate hydrogel transfer layer. (a) Steps for the Au NIL-array transfer to alginate hydrogel. (i) First, we transferred the Au NIL-array to a glass coverslip and then functionalized the Au with cysteamine. (ii) Next, we cast the alginate hydrogel transfer layer on top of the Au NIL-array. (iii) Then, we peeled off the alginate hydrogel containing the Au NIL-array and placed it pattern-side up. (b) Optical image of the Au NIL-array printed alginate hydrogel. (c-d) Top view SEM images of the Au (c) NIL-dot and (d) NIL-wire printed alginate hydrogel transfer layers.

## Cell culture on Au NIL-array printed alginate hydrogels

We investigated the viability, migration, and morphology of embryonic mouse fibroblast cells (NIH/3T3-GFP) on two different Au nanopatterns: dots (approximately 250 nm diameter, 550 nm center-to-center spacing, 300 nm rim-to-rim spacing) and wires (approximately 300 nm wide, 450 nm spacing). As mammalian cells are known to have poor adhesion to alginate hydrogels, ^30^ we bioconjugated gelatin to the Au NIL-array printed hydrogels prior to seeding NIH/3T3-GFP cells. The bioconjugation process (Figure S4) involves sequential functionalization of the Au surface with cysteamine and glutaraldehyde, and subsequent coating with 0.1% gelatin (Bloom 300, Type A). Glutaraldehyde contains aldehyde groups at both ends of the molecule and can bind to the amine groups in cysteamine and gelatin.^31^

Of note, certain studies have reported that glutaraldehyde can undergo acetalization with the hydroxyl groups of alginate.^32^ However, this crosslinking reaction occurs only under acidic conditions.^33^ Conversely, glutaraldehyde exhibits rapid reactivity with amine groups and forms thermally and chemically stable crosslinks around neutral pH.^34^ Therefore, the reaction conditions, including pH, concentration, temperature, and reaction times, must be optimized to achieve the desired crosslinking of glutaraldehyde with cysteamine and gelatin, instead of alginate.^35^ Approximately 24 hours after seeding the cells on the Au NIL-array printed hydrogels, we observed that the NIH/3T3-GFP cells on the NIL-wire printed hydrogel preferably migrated parallel to the nanowires, whereas those on NIL-dots exhibited random migration (Figure 2a-c). Using ImageJ, we estimated the elongation factor of the fibroblasts on the Au NIL-array printed hydrogels by measuring the long axis length over the short axis length of the cell. The elongation factor of the cells on the Au NIL-wire printed hydrogel was approximately twice that of the cells on the Au NIL-dot printed hydrogel (Figure 2d). This observation suggests that the gelatin selectively conjugated to the Au NIL-arrays, thus enhancing cell alignment and elongation on the Au NIL-wire printed alginate hydrogel compared to the Au NIL-dot printed alginate hydrogel. Also, cells on the Au NIL-dot printed hydrogel migrated about 1.4 times faster than cells on the Au NIL-wire printed hydrogel (Figure 2e). We note that the relationship between surface patterning and cell migration speed is nuanced and dependent on a variety of factors, such as relative adhesion, relative modulus, and pattern dimension and density.^36-39^ While many prior studies on cell migration have been carried out on stiffer substrates like glass or polydimethylsiloxane (PDMS), our method enables fabrication of soft hydrogel and anatomically relevant substrates with tunable and precisely engineered surface patterns for investigation cell morphology and dynamics.

**Figure 2.**
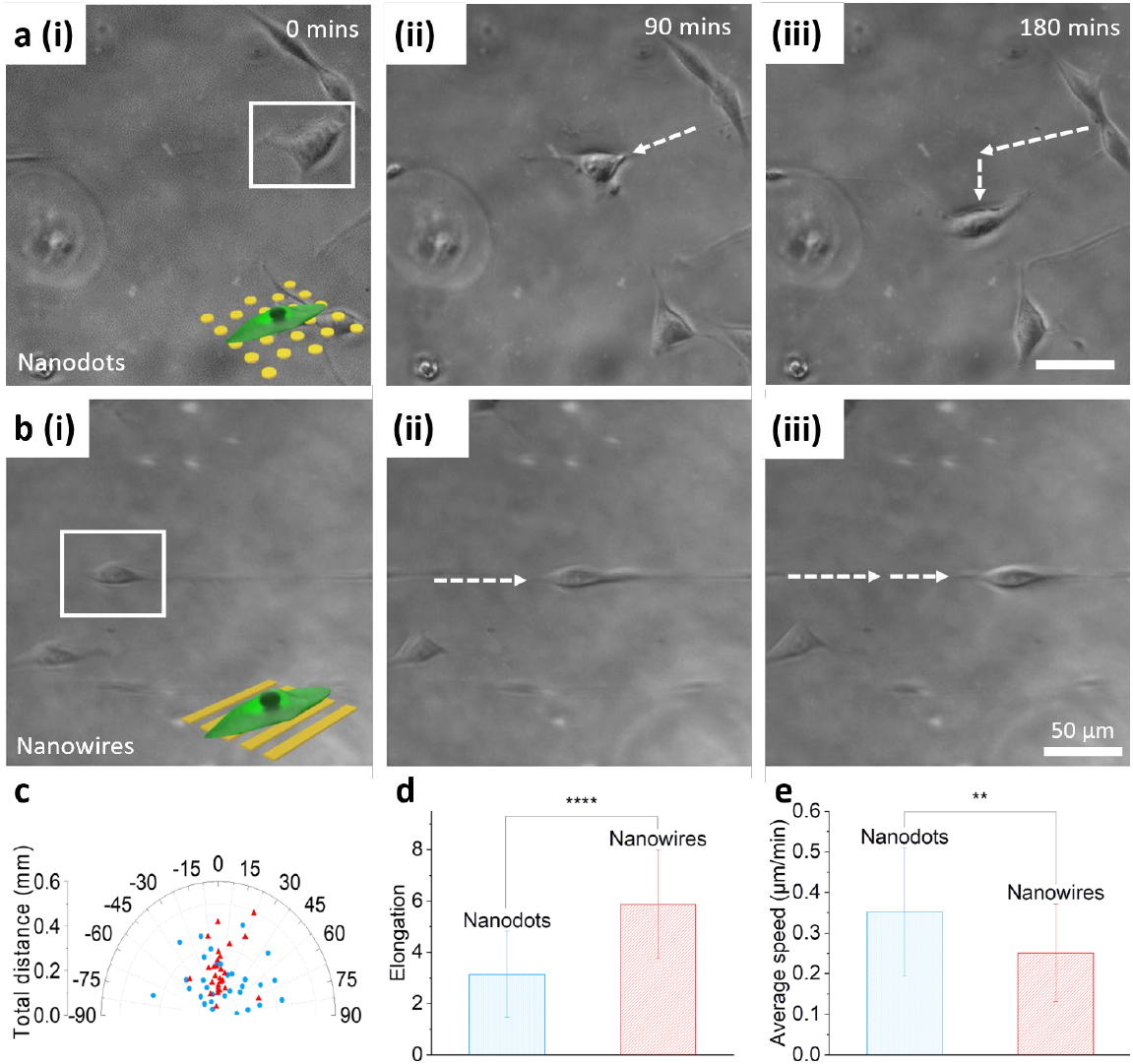
Characterization of NIH/3T3-GFP fibroblast migration on the Au NIL-array printed alginate hydrogels. (a) Time-lapse confocal phase images of a representative NIH/3T3-GFP cell on the Au NIL-dot printed alginate hydrogel show changing migration trajectory. (b) Time-lapse confocal phase images of a representative NIH/3T3-GFP cell on the Au NIL-wire printed alginate hydrogel show linear migration trajectory. (c) Angular distribution plot of the cell migration orientation and distance. The plot shows that cells migrated on the Au NIL-dots (blue) with no significant directional preference, whereas cells on the Au NIL-wires (red) migrated primarily along the direction of the wires. (d) Elongation factor of the cells on Au NIL-dots (blue) and Au NIL-wires (red). Cell elongation is more pronounced on Au NIL-wires. Data are presented as mean ±SD (n = 30 cells). Statistical analysis was performed using the unpaired two-sided t test. ****p<0.0001. (e) The average cell migration speed is higher on the Au NIL-dots (blue) than on the Au NIL-wires (red). Data are presented as mean ±SD (n = 30 cells). Statistical analysis was performed using the unpaired two-sided t test. **p<0.01.

## Biotransfer printing Au NIL-arrays on rat brains

Alginate hydrogel is not only biocompatible with cells and tissues but can also undergo reverse gelation by metal chelates (e.g., EDTA) and specific enzymes (e.g., alginate lyases).^40^ Hence, it is an attractive sacrificial material for transferring the Au NIL-arrays onto living organs and cells. To demonstrate, we biotransfer printed Au NIL-arrays on a whole rat brain (Figure 3b-e) and on a rat brain slice (Figure S5 a). First, we dissected brains from 21-day postnatal rats and positioned the Au NIL-wire printed alginate hydrogels on the cerebral cortex of a whole brain and on a coronal brain slice. After leaving the samples in cell culture media for 2 hours (Figure 3a), we dissociated the alginate hydrogels with 20 mM EDTA. We observed that the Au NIL-wires remained bonded to the surface of the whole brain (Figure 3 d-f). In contrast, the Au NIL-wires on the coronal brain slice did not adhere and were washed away after rinsing with EDTA (Figure S5 b). Prior studies have shown that thin film patterns can conform to brain surfaces through physical adhesion forces like water capillarity or by interfacial hydrogel layers.^41-44^ In our experiments, the Au NIL-arrays selectively adhered to the surface of the whole rat brain, which exhibits distinct cell and matrix compositions from the coronal brain slice. This suggests that the Au NIL-array adhesion mechanism may be cell-type specific and cell adhesion related. Further studies are needed to determine the specific adhesion factors on different cell and tissue interfaces.

**Figure 3.**
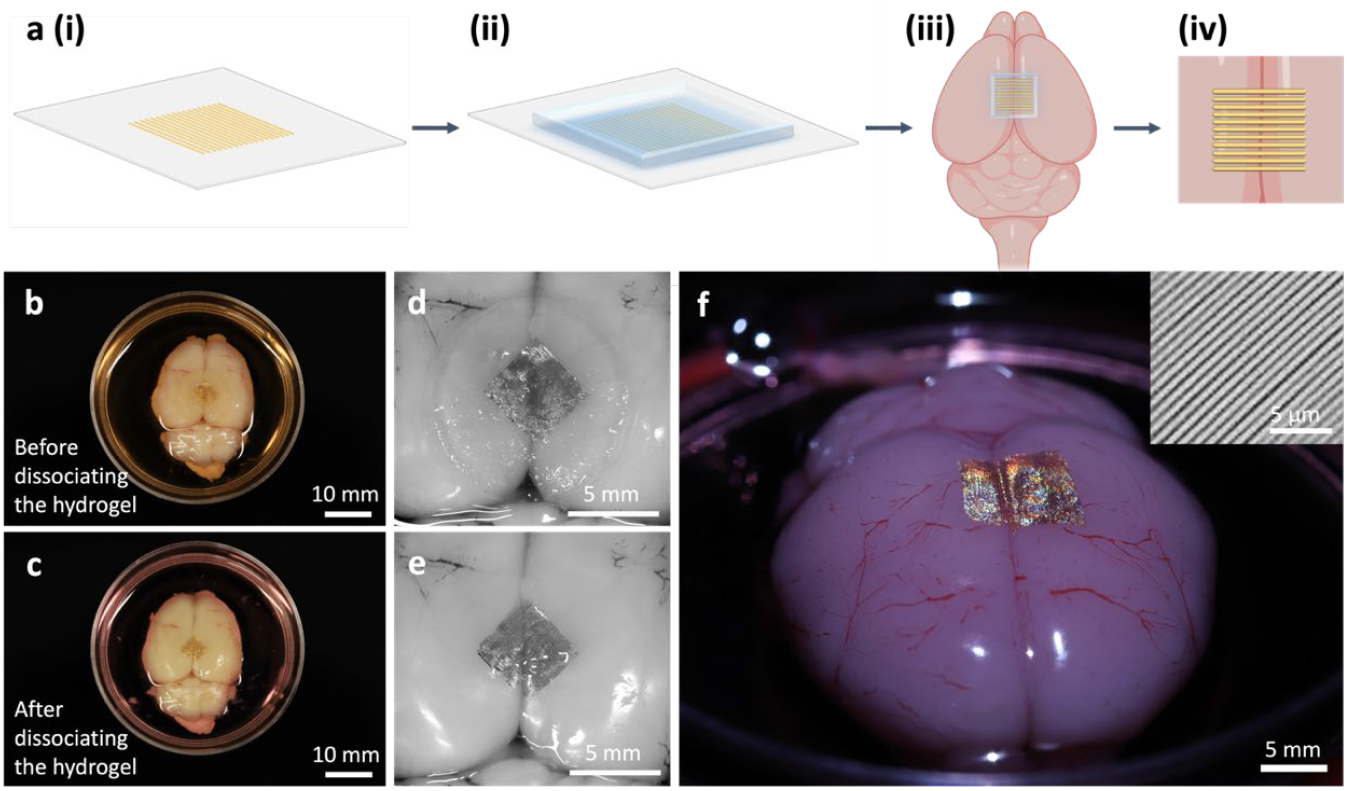
Schematic illustration and microscopy images showing the biotransfer printing process of Au NIL-wires on rat brains. (a) Steps for the Au NIL-array transfer to a rat brain. (i) First, we transferred the Au NIL-array to a glass coverslip and functionalized the Au surface with cysteamine. (ii) Next, we cast alginate hydrogel on top of the Au NIL-array. (iii) Then, we peeled off the alginate hydrogel containing the Au NIL-array and bioconjugated the patterned surface with gelatin before placing it on top of a rat brain. (iv) Finally, we dissociated the alginate hydrogel with 20 mM EDTA. (b-c) Optical and (d-e) magnified images of Au NIL-wires on a rat brain (b, d) before and (c, e) after dissociation of the alginate hydrogel with 20 mM EDTA. (f) Side view image of Au NIL-wires on a rat brain after dissociation of the alginate hydrogel transfer layer with 20 mM EDTA. The inset shows a magnified view of the Au NIL-wires on the rat brain obtained by a laser scanning microscope.

## Biotransfer printing Au NIL-arrays on live NIH/3T3-GFP cell sheets

To assess the biotransfer printing capacity at the single-cell level, we transferred the Au NIL-arrays onto a monolayer of fibroblasts. The biotransfer printing process demonstrated here is inspired by cell sheet transfer with slight modifications (Figure 4a). Briefly, we cultured NIH/3T3-GFP monolayer cell sheets on Au NIL-array printed alginate hydrogels for about 24 hours. Then we flipped over the cell-seeded hydrogels onto gelatin-coated coverslips and let the cells attach to the coverslips overnight. We dissociated the alginate hydrogels by rinsing them with 20 mM EDTA for about 9 minutes. After dissociating the alginate hydrogels, we analyzed the viability of the NIH/3T3-GFP cell sheets with Au NIL-arrays using propidium iodide (PI). Using fluorescence microscopy, we qualitatively observed high cell viability with both Au NIL-dots and NIL-wires (Figure 4b-c). Specifically, the fibroblasts patterned with Au NIL-dots had a viability of approximately 97%, while those patterned with Au NIL-wires had a viability of approximately 98% (Figure 4d). These results indicate that this transfer printing process is biocompatible with live cells. We also observed reflective colors from the fibroblast cell sheet patterned with the Au NIL-array (Figure 4e), which suggests that the shape of the nanopattern array was retained on the cell sheet. We fixed the cell sheets patterned with Au NIL-arrays for SEM analysis and noticed that both the Au NIL-dots (Figure 4f) and NIL-wires (Figure 4g) achieved conformal contact with the cell sheets. However, when the Au NIL-arrays were biotransfer printed on cells cultured for shorter times, such as 4 hours, the patterns were either distorted or fragmented (Figure S6 b). Concurrently, we observed the presence of a thin, porous film on the cells cultured for at least 24 hours (Figure S7) but not on the cells cultured for a shorter period (Figure S6 a-b). Based on these SEM images, we infer that this thin, porous film represents extracellular matrix (ECM) secreted by NIH/3T3-GFP cells and that it may be involved in facilitating adhesion between the cell sheets and the Au NIL-arrays.^45,46^

**Figure 4.**
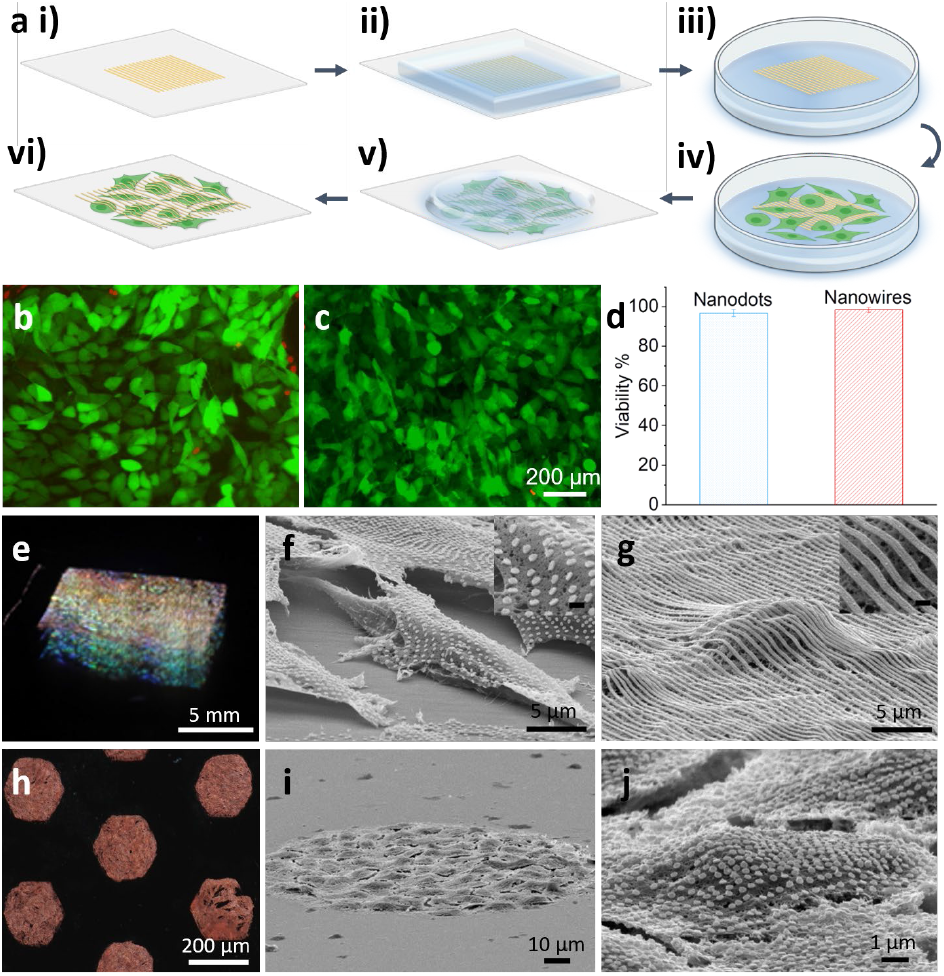
Characterization of the biotransfer printed Au NIL-arrays on NIH/3T3-GFP cell sheets. (a) Schematic illustration showing the steps for transferring the Au NIL-array to a cell sheet. (i) First, we transferred the Au NIL-array to a glass coverslip and functionalized the Au with cysteamine. (ii) Next, we cast alginate hydrogel on top of the Au NIL-array. (iii) Then we peeled off the alginate hydrogel containing the Au NIL-array and bioconjugated the patterned surface with gelatin. (iv) Then we seeded NIH/3T3-GFP cells on top of the Au NIL-array printed hydrogel. (v) Next, we flipped over the cell-seeded hydrogel onto a gelatin-coated coverslip. (vi) Finally, we dissociated the alginate hydrogel with 20 mM EDTA. (b-c) Live-dead assay of NIH/3T3-GFP cells patterned with Au (b) NIL-dots or (c) NIL-wires after dissociating alginate hydrogel with 20 mM EDTA. Dead cells in red fluorescence. (d) Viability of the cells with Au NIL-dots or NIL-wires. (e) Optical image of an 8 × 8 mm array of Au NIL-arrays printed on an NIH/3T3-GFP cell sheet. (f) SEM image of Au NIL-dots on fibroblasts. (g) SEM image of Au NIL-wires on fibroblasts. (h) Optical image of the micro-cell patches with Au NIL-dots. (i) SEM image of a micro-cell patch with Au NIL-dots. (j) Magnified SEM image of a cell in the micro-cell patch with Au NIL-dots.

Our fabrication process is not only compatible with NIL but also with microscale photolithography (Figure 4h-j). To illustrate this feature, we used photolithography and wet etching to define 200 μm-wide hexagonal patches and 200 μm-wide triangular patches of Au NIL-arrays. We then biotransfer printed the micro-patches of Au NIL-arrays onto cell sheets according to the steps shown in Figure 4a. The bioconjugation of gelatin to the Au surface resulted in the selective growth of fibroblast cells on the micro-patches of Au NIL-arrays, as the alginate hydrogel itself does not contain any cell-adhesion ligands necessary for promoting cell attachment (Figure 4h). After dissociating the alginate hydrogel with EDTA, we observed a high yield of Au NIL-array printed micro-cell patches attached to the coverslips. The supplementary movies (Movie S1-3) show the migration of cells with patches of Au NIL-wires biotransfer printed on top of the cells. The cells with Au NIL-wires appear healthy and able to migrate indicating biocompatibility of the transfer process. Also, the movies provide evidence that the Au NIL-wires can adhere and move with cells during this 16 hr period of migration.

In summary, we have introduced a new approach for creating nano-bio interfaces in the form of Au NIL-arrays on live cells and tissues. Our approach utilizes molecular linkers for the careful manipulation of adhesion between materials and alginate hydrogel as both a cell culture scaffold and degradable transfer layer. We have demonstrated the ability of the Au NIL-arrays printed on anatomically relevant and ultrasoft substrates, such as hydrogels, to guide cell orientation and migration. By dissociating the alginate hydrogel with EDTA, we achieved conformal contact between the Au NIL-arrays and *ex vivo* rat brains as well as live cells. Noting the variation in adhesion strength among different cell types and culture methods, additional studies are needed to characterize and optimize the specific adhesion mechanisms for robust long-term bonding. Importantly, NIL patterning enables facile integration of multifunctional devices in a high throughput manner.^47–49^ Therefore, this approach could enable advanced functional optical and electronic devices, such as metamaterial arrays, plasmonic sensors, transistors, circuits, and antennas, to be imprinted on hydrogels, live cells and tissues.^19,22,50-52^ We expect this nanopatterning process, combined with various classes of materials and standard microfabrication techniques like photolithography and e-beam lithography, to open opportunities for the development of new cell culture substrates, biohybrid materials, bionic devices, and biosensors.

## Acknowledgments

We acknowledge support from the Air Force Office of Scientific Research 21RT0264– FA9550-21-1-0284. Research reported in this publication was partially supported by the National Institute on Aging of the National Institutes of Health under Award Number R03AG073834. The content is solely the responsibility of the authors and does not necessarily represent the official views of the National Institutes of Health.

