## Supplementary Information for "Toward single cell tattoos: Biotransfer printing of lithographic gold nanopatterns on live cells"

**Author Contributions**

‡These authors contributed equally.

**Fabrication of the Au NIL-arrays on glass coverslips**

We fabricated the Au nanopattern arrays via nanoimprint lithography (NIL). Briefly, we spin-coated a layer of polymethylglutarimide (PMGI SF6, Kayaku Advanced Materials) as the sacrificial layer on a silicon (Si) wafer. Then we spin-coated a layer of NIL resist (mr-I 7030, Micro Resist Technology) with a thickness of 350 nm and thermally imprinted the nanopatterns using a Nanonex Advanced Nanoimprint Tool NX-B200 with a pressure of 350 psi at 130 °C. We used a commercial, low-cost Si master stamp (LightSmyth grating) of two kinds to create nanodots (approximately 250 nm diameter, 550 nm center-to-center spacing, 300 nm rim-to-rim spacing) and nanowires (approximately 300 nm width, 450 nm spacing). After imprinting, we etched away the residual NIL resist with oxygen plasma at 60 W for 2 minutes. We used thermal evaporation to deposit 50 nm of Au with a 5 nm-thick Cr layer underneath to improve adhesion to the nanopatterned wafer. After deposition, we sonicated the sample in acetone to completely dissolve the NIL resist and obtained a large-area array (8 by 8 mm) of Au nanopatterns on the Si wafer.

We spin-coated a layer of polymethyl methacrylate (PMMA A4) on top of the Au NIL-array as a carrier film. We released the Au NIL-array from the Si wafer by dissolving the PMGI sacrificial layer in a positive photoresist developer (MF-26A). To retain the shape of the nanopatterns, it is important to keep the thin film floating on the surface of the liquid for these steps. We rinsed the film with water by displacing the photoresist developer with deionized (DI) water three times. Then we etched the Cr in Cr etchant (Cr Cermet Etchant TFE, Transene) and repeated the rinsing step with water. Afterward, we picked up the film from the water-air interface using a glass coverslip. After air-drying the film, we etched the PMMA film in oxygen plasma at 60 W for 30 minutes.

**Alginate hydrogel preparation**

We used sodium alginate with high guluronic acid block content (average MW 177kDa, I1G, KIMICA) to fabricate the alginate hydrogel based on a published method.[1] Briefly, we purified alginate by dialyzing it against DI water for 3 days with a 3500 MWCO membrane. Afterward, we used activated charcoal and sterile filtration to purify the alginate. We lyophilized the purified alginate for 4-5 days and stored it at -20℃ until needed.

**Cell culture**

We cultured the NIH/3T3-GFP cells (kindly provided by Dr. Yun Chen at Johns Hopkins University) in standard DMEM (Gibco) with 10% fetal bovine serum (HyClone) and 1% penicillin/streptomycin (Gibco). The cells were cultured in a humidified incubator at 37 °C with 5% $CO_2$, kept at sub-confluency, and passaged every 2-3 days.

### Transfer of the Au NIL-arrays to cell sheets and rat brains

We transferred the Au NIL-arrays from the glass coverslips onto cell sheets and tissues using alginate hydrogel as a biocompatible and sacrificial transfer layer. To facilitate the delamination of the Au NIL-array from the glass coverslip, we enhanced relative adhesion to the alginate hydrogel by chemically modifying the Au surface with a self-assembled monolayer of cysteamine. We immersed the Au NIL-array in a 0.26 mM cysteamine ethanol solution for an hour. Then we prepared the alginate hydrogel by mixing 0.5 ml of the 2.5 wt% alginate solution with 125 µl of 25 mM calcium sulfate. We obtained a homogenous mixture by loading each solution in a syringe and mixing with a dual Luer-lock connector. Then we cast the alginate hydrogel on the Au NIL-array and allowed the solution to gel for 45 minutes under a glass slide with 1 mm-thick spacers. Afterward, we carefully peeled off the alginate hydrogel containing the Au NIL-array from the glass coverslip and placed it pattern-side up in a petri dish. We sterilized the hydrogel by placing it under UV light for an hour and made sure to immerse it in excess $CaCl_2$ solution to prevent dehydration. Then we repeated the cysteamine functionalization step and rinsed the alginate hydrogel three times with DI water. To bind the gelatin molecules, we immersed the Au NIL-array in a 17.6 mM glutaraldehyde water solution for 30 minutes and rinsed the hydrogel three times with DI water. Next, we immersed the hydrogel in a 0.1% gelatin (Bloom 300, Type A) phosphate-buffered saline solution for an hour and aspirated the excess solution. We used the same gelatin coating procedure to obtain the gelatin-coated glass coverslips. After seeding NIH/3T3-GFP cells on the Au NIL-array printed alginate hydrogel, we placed it in an incubator for 24 hours. To obtain the Au NIL-array printed cells, we picked up the cell-seeded hydrogel and flipped it over onto a gelatin-coated coverslip so that the cells were in direct contact with the gelatin-coated coverslip. We allowed the cells to attach to the gelatin-coated coverslip overnight and finally dissociated the alginate hydrogel by rinsing it with 20 mM of EDTA for about 9 minutes. For the transfer of the Au NIL-arrays to rat brains, we repeated the same alginate hydrogel casting and gelatin conjugation steps. Then we placed the Au NIL-array printed alginate hydrogel on top of the brain tissue so that the Au NIL-array was in direct contact with the tissue surface. After leaving the samples in cell culture media for about 2 hours, we dissociated the alginate hydrogel by rinsing it with 20 mM EDTA for about 9 minutes.

### Cell tracking and imaging

We used a Nikon TE2000 microscope with 10X objective lens to capture cell movement over 14 hours at 5-minute intervals. During imaging, cells were maintained on a temperature and $CO_2$-controlled stage in an incubator at 37 °C and 5% $CO_2$. We used CellTracker software to record cell migration paths and calculate cell migration speed.[2]

The inset in Figure 3f was obtained using a Keyence laser scanning microscope VK-X100.

**Animal experiments**

Sprague Dawley rats were purchased from Charles River and Taconic Biosciences, and the animals were bred and housed at Johns Hopkins animal facilities. All animal procedures and experiments were performed in accordance with guidelines set by the National Institutes of Health and the Johns Hopkins University Animal Care and Use Committee (ACUC). Postnatal 21-day rats were euthanized using carbon dioxide ($CO_2$). Rats were further subjected to cervical dislocation following euthanasia by $CO_2$ inhalation. Decapitation was performed, and brains were dissected for follow-up experiments.

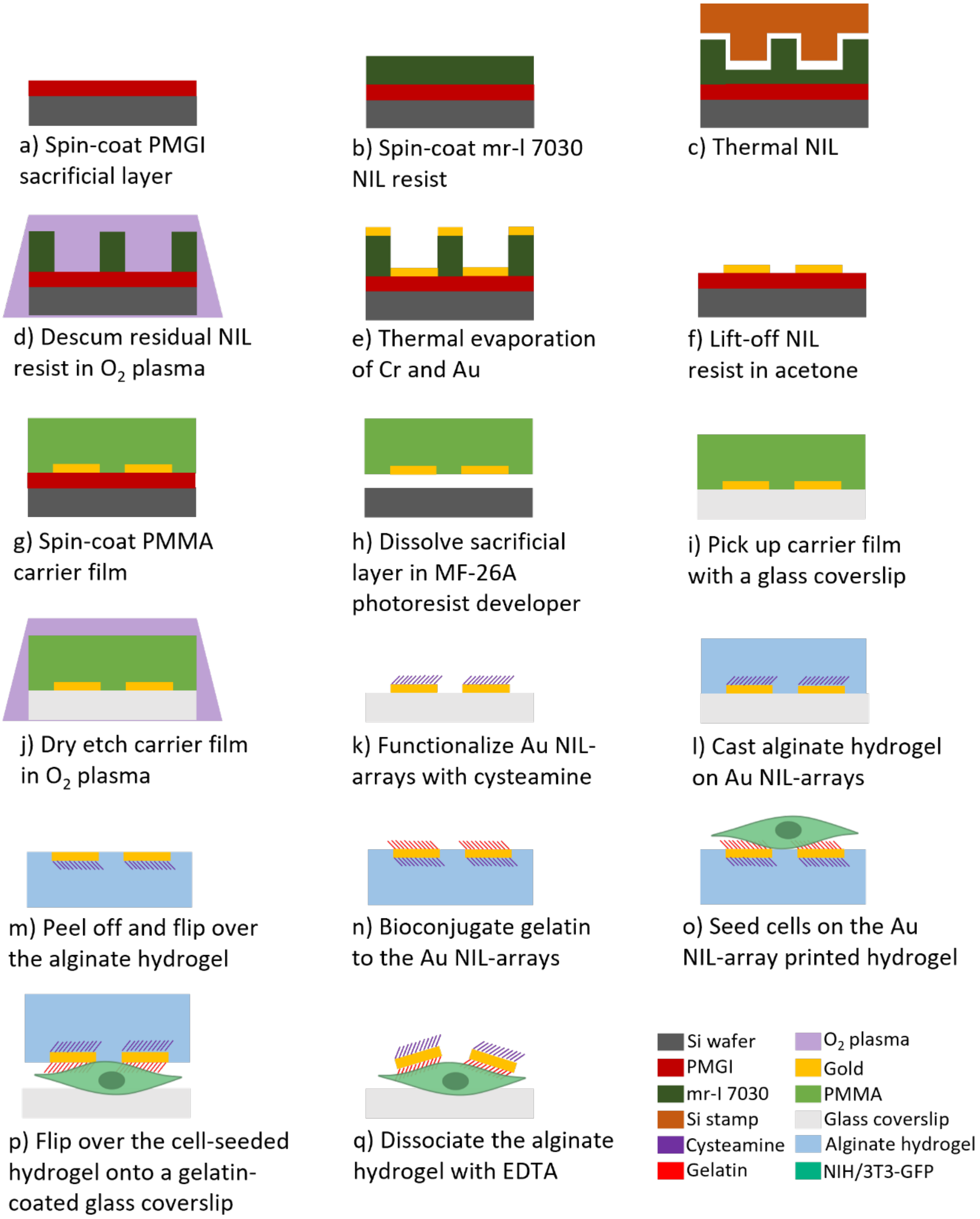


**Figure S1.** Schematic illustration showing the step-by-step fabrication of the Au NIL-arrays on Si wafers and subsequent transfer onto glass coverslips and cell sheets. The main steps include (a) spin-coating the sacrificial layer (PMGI) on a Si wafer, (b) spin-coating the NIL resist (mr-I 7030), (c) thermal NIL using a Si stamp, (d) descumming the residual NIL resist with oxygen plasma, (e) thermally evaporating 5 nm of Cr and 50 nm of Au, (f) lifting off the NIL resist in acetone, (g) spin-coating the carrier film (PMMA), (h) dissolving the sacrificial layer in positive photoresist developer (MF-26A) and Cr in Cr etchant (Cr Cermet Etchant TFE) followed by rinsing with water, (i) manually picking up the Au NIL-array from the water-air interface using a glass coverslip, (j) removing the carrier film using an oxygen plasma, (k) functionalizing the Au NIL-array with cysteamine, l) casting alginate hydrogel on the Au NIL-array, m) peeling off the Au Nil-array printed alginate hydrogel from the glass coverslip and placing it pattern-side up, n) bioconjugating gelatin to the Au NIL-array according to the process detailed in Figure S4, o) seeding NIH/3T3-GFP cells on the Au NIL-array printed hydrogel and culturing for 24 hours, p) flipping over the cell-seeded hydrogel onto a gelatin-coated glass coverslip, and q) dissociating the alginate hydrogel with 20 mM EDTA to obtain the Au NIL-array printed cells.

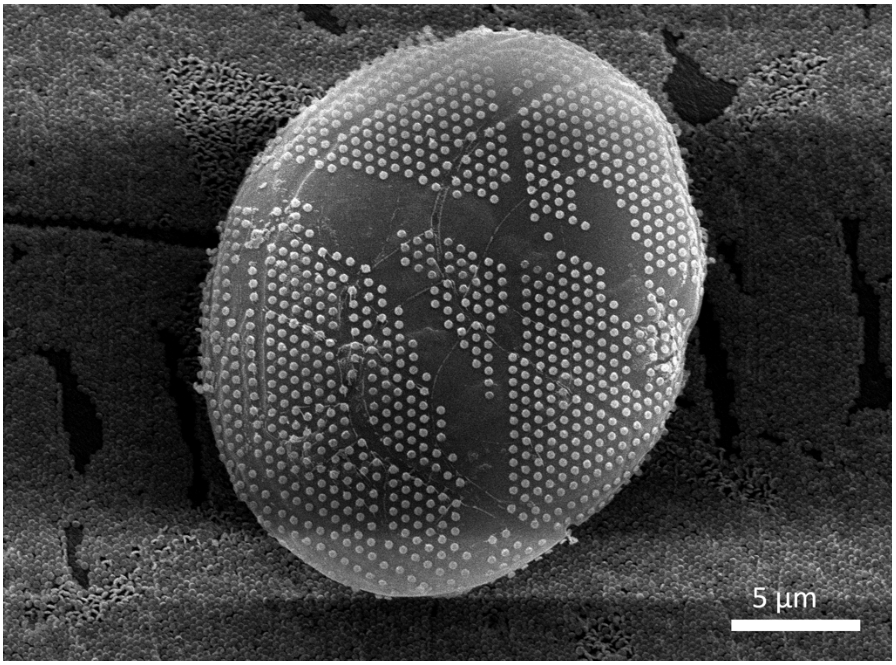


**Figure S2.** SEM image of Au NIL-dots bioitransfer printed on the surface of a 3D microparticle.

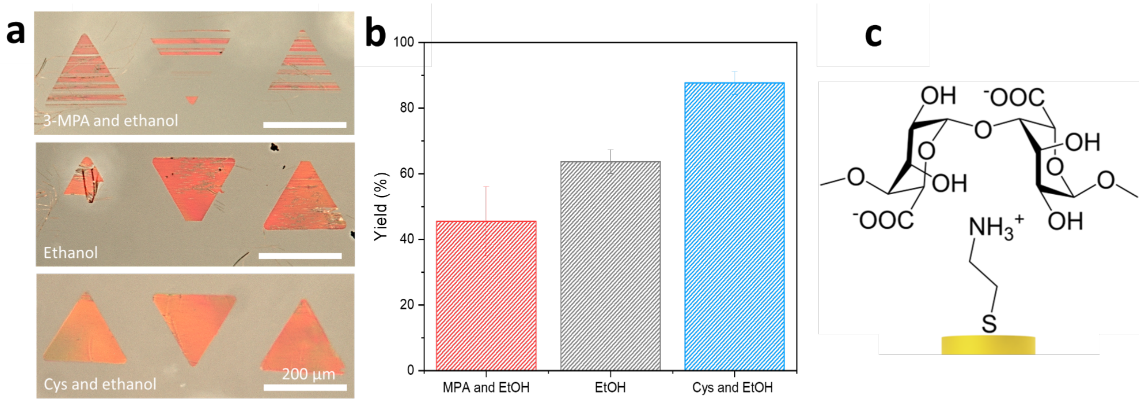


**Figure S3.** Characterization of the Au NIL-arrays on alginate hydrogels. a) Optical images of the Au NIL-arrays functionalized with 3-mercaptopropionic acid and ethanol (top), ethanol (middle), or cysteamine and ethanol (bottom). b) Transfer yield of the Au NIL-arrays functionalized with different molecules from the glass coverslips to the alginate hydrogels. c) Proposed mechanism for enhancing the transfer yield of the Au NIL-arrays to the alginate hydrogel with positively charged cysteamine.

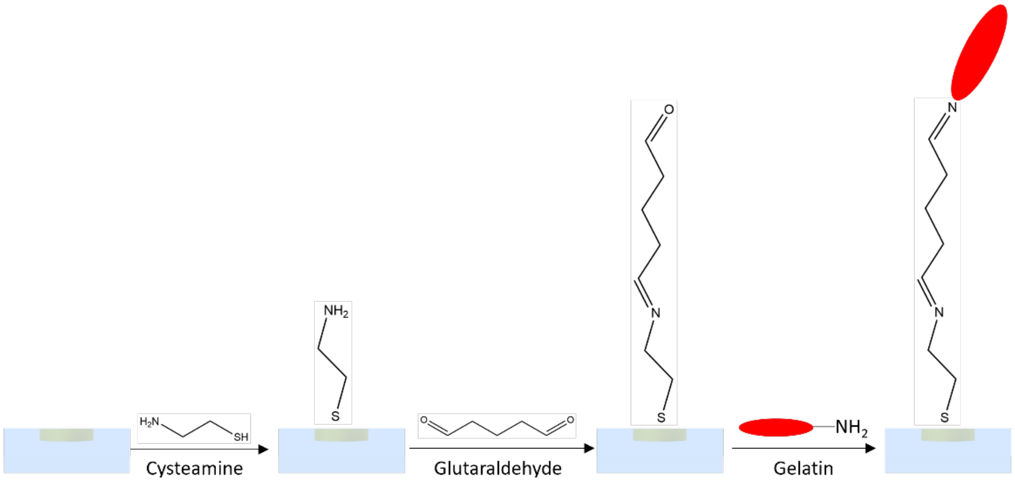


**Figure S4**. Schematic illustration showing the steps for the bioconjugation of gelatin to the Au NIL-arrays.

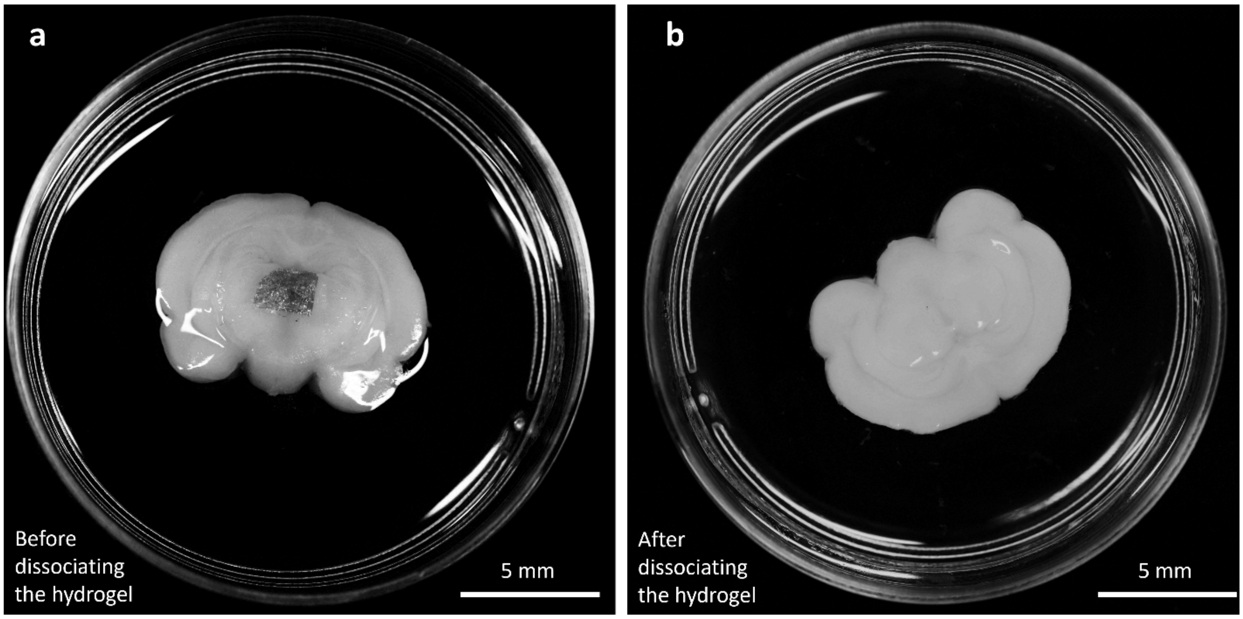


**Figure S5.** Characterization of the biotransfer printed Au NIL-wires on a rat brain slice. (a) Optical image of the Au NIL-wires on a rat brain slice before dissociating the alginate hydrogel. (b) Optical image of the Au NIL-wires on a rat brain slice after dissociating the alginate hydrogel with 20 mM EDTA.

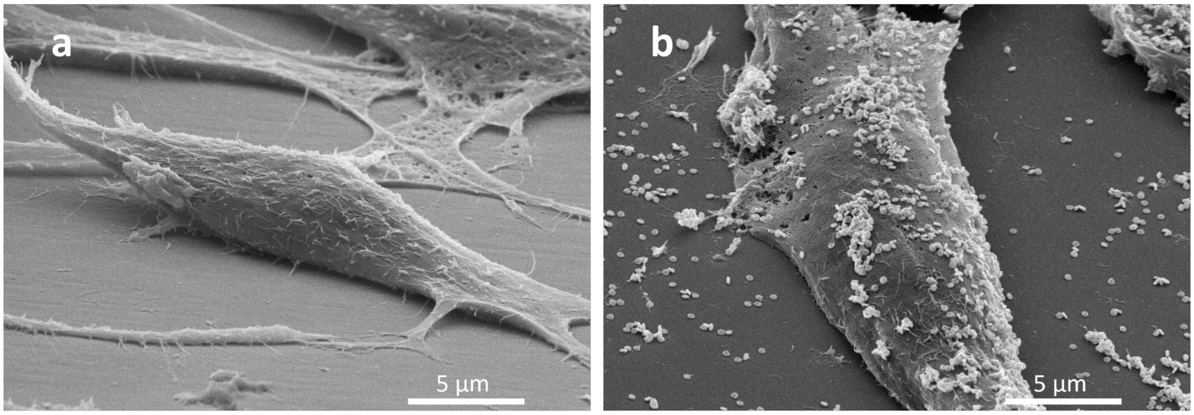


**Figure S6**. SEM images of NIH/3T3-GFP fibroblasts. a) SEM image of cells without Au NIL-arrays on a glass coverslip, b) SEM image of distorted Au NIL-dots on cells that were cultured for 4 hours.

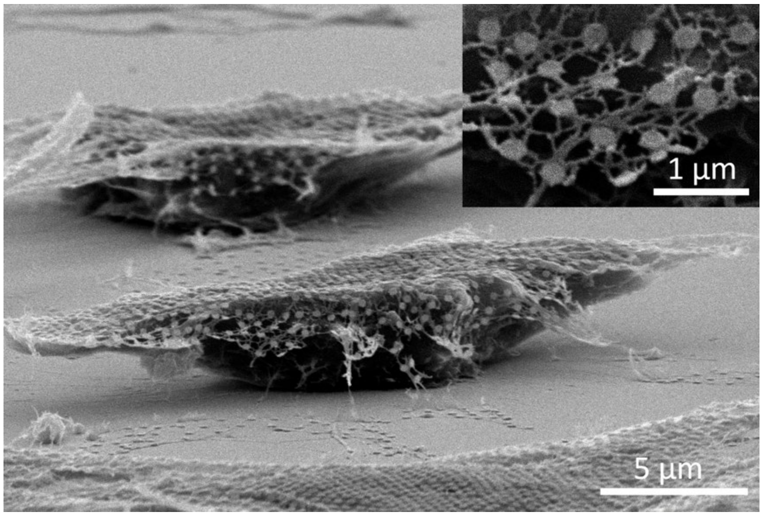


**Figure S7**. SEM image of NIH/3T3-GFP fibroblasts covered with a thin, porous film and Au NIL-dots.

**Supplementary movie S1:** Zoomed-in video clip of live cell imaging assembled from images captured over a period of 16.1 hours at 5-minute intervals with the Nikon TE2000 microscope. The clip shows an NIH/3T3-GFP cell with Au NIL-wires biotransfer printed on top. The cell appears healthy and able to migrate indicating biocompatibility of the transfer process, and the Au NIL-wires move with the cell during migration over this 16 hr period.

**Supplementary movie S2:** Zoomed-in video clip of live cell imaging assembled from images captured over a period of 16.1 hours at 5-minute intervals with the Nikon TE2000 microscope. The clip shows a different NIH/3T3-GFP cell with Au NIL-wires biotransfer printed on top. The cell appears healthy and able to migrate indicating biocompatibility of the transfer process, and the Au NIL-wires move with the cell during migration over this 16 hr period.

**Supplementary movie S3:** Large area video clip of live cell imaging assembled from images captured over a period of 16.1 hours at 5-minute intervals using the Nikon TE2000 microscope. The clip shows NIH/3T3-GFP cells with Au NIL-wires biotransfer printed on top. The cells with NIL-wires appear healthy and able to migrate indicating biocompatibility of the transfer process. Au NIL-wires adhere to some cells and move with them during migration over this 16 hr period. The images were taken with a 10X objective lens.
